# Cargo and Cell Specific Differences in Extracellular Vesicle Populations Identified by Multiplexed Immunofluorescent Analysis

**DOI:** 10.1101/735936

**Authors:** Kevin Burbidge, Virginia Zwikelmaier, Benjamin Cook, Grace Ispas, David J. Rademacher, Edward M. Campbell

## Abstract

Extracellular vesicles (EVs) are implicated in a wide variety of biological activities, have been implicated in the pathogenesis of numerous diseases, and have been proposed to serve as potential biomarkers of disease in human patients and animal models. However, characterization of EV populations is often performed using methods that do not account for heterogeneity of EV populations and require comparatively large sample sizes to facilitate analysis. Here, we describe an imaging-based method that allows for the multiplexed characterization of EV populations at the single EV level following centrifugation of EV populations directly onto cover slips, allowing comprehensive analysis of EV populations with relatively small samples. We observe that canonical EV markers are present on subsets of EVs which differ substantially in a producer cell and cargo specific fashion, including differences in EVs containing different HIV-1 proteins previously reported to be incorporated into pathogenic EVs. We also describe a lectin binding assay to interrogate EVs based on their glycan content, which we observe to change in response to pharmacological modulation of secretory autophagy pathways. These studies collectively reveal that multiplexed analysis of EV populations using fluorescent microscopy can reveal differences in specific EV populations that may be used to understand the biogenesis of specific EV populations or to interrogate small subsets of EVs of interest within larger EV populations.

## Introduction

Cells release populations of extracellular vesicles (EVs) of various sizes which contain a multitude of cargoes including RNA, proteins, and lipids(1-3). While EVs were once thought to be relegated strictly to the task of shedding unwanted cellular waste, it is now appreciated that EVs play an integral role in cell-to-cell communication(4) and that all cell types produce EVs under both healthy and pathological cellular states(5). As a result, understanding mechanisms associated with the transport and characterization of EVs has become an increasingly popular focus. In particular, by identifying specific EV cargoes under pathological conditions, EVs have the potential to be excellent biomarkers to assist in earlier detection and diagnosis of a variety of diseases(6, 7).

The most often cited subtypes of EVs are exosomes, microvesicles, and apoptotic bodies, which are derived from distinct mechanisms and sub-cellular compartments. As a result, the different EV subtypes are characterized based on their size, protein composition and origin(3, 8-10). Exosomes are typically appreciated to range between 30-150nm and originate as intraluminal vesicles (ILVs) formed within multivesicular bodies (MVBs)(3, 8-10). The process of secretory autophagy leads to the release of an EV population similar to exosomes, which is known to be upregulated in the context of lysosomal dysfunction(11-14). In comparison, microvesicles have a more dynamic size range at 20-1000nm and bud directly from the plasma-membrane, while apoptotic bodies are much larger and thought to be over 1000nm in diameter and originate from dying cells(3, 8-10). In addition to size, researchers often validate the isolation of exosomes or EVs by demonstrating the presence of specific cellular proteins, including the tetraspanins CD9, CD63, and CD81(1, 8, 15, 16).

Experimentally, EV populations are typically isolated and separated from cell debris via a series of centrifugation steps(17). Other commonly employed isolation methods include filtration and purification based on highly enriched EV markers. These isolated EVs can then be used for a variety of assays, such as western blots, ELISAs, or mass spectrometry to measure protein composition. Utilizing these techniques and defining EVs by these previously stated criterion has led to an impressive advancement in understanding the vast differences in EV composition and potential impact on neighboring cells(18). However, there is also accumulating evidence indicating that in spite of existing criteria to classify EVs, heterogeneous populations of EVs exist(8-10, 18, 19) and these subpopulations of EVs are insufficiently resolved using currently used approaches. Other studies reveal that alterations of cellular homeostasis in EV producing cell can alter the quantity as well as the quality of the EVs released from cells observation that EVs released from cells (20). These observations suggest that bulk analysis of EV populations following centrifugation or other methods that analyze the molecular composition of all EVs in a sample fail to reveal heterogeneity in EV populations, especially in cases where potentially pathological or therapeutic populations of EVs exist as a subpopulation within a larger population of EVs. Thus, it is increasingly important that methods be created to assist in distinguishing EVs without the need to disrupt EV heterogeneity.

To address this issue, we formulated an imaging-based workflowthatutilizes quantitative fluorescent microscopy to characterize individual <u>EV</u>s via <u>M</u>ultiplexed <u>A</u>nalysis of <u>C</u>o-localization (EV-MAC) of their protein and glycan determinants. This method allows populations of EVs, and EVs containing specific cargos, to be comparatively analyzed for the presence or absence of other cellular markers known to be present on normal or pathological EV populations. To validate this methodology, we characterized EVs produced from cells expressing HIV-1 proteins which are reported to be extruded in pathological exosomes(21) and observed differences in the degree to which individual cellular markers associated with EVs containing each cargo. We also examined the EVs released from cells following inhibition of lysosomal degradation, which is known to induce the release of pathological EVs in the context of neurodegenerative disease (22, 23) and observe that lysosomal inhibition alters the glycan signature of the EV population released from cells under these conditions. Lastly, we applied this technique to human biological fluids from healthy donors, examining EVs isolated from saliva and plasma. Collectively, these results demonstrate that multiplexed analysis of EV populations can be used to interrogate the protein or glycan content of specific sub-populations of EVs of interest, which could potentially provide insight into the biogenesis of EV populations or aid in the discovery of biomarkers associated with pathological EVs.

## Materials and Methods

### Cell Culture

The HEK293T (CRL-3216) and the THP-1 (TIB-202) immortalized cell-lines were purchased from the American Type Culture Collection (ATCC). HEK293T and THP-1 cells were cultured in an incubator at 37C and 5% CO2 in Dulbecco’s modified Eagle’s Medium (DMEM) containing phenol red (Invitrogen) or Rosewell Parks Memorial Institute (RPMI) 1640, supplemented with the 10% fetal bovine serum (FBS) (Hyclone), 10ug/ml ciprofloxacin hydrochloride, 100IU/ml penicillin, and 100ug/ml streptomycin.

### Stabile Expression of S15 mCherry Construct

HEK293Tand THP-1 cells were transduced to stably express S15-mCherry using the lentiviral vector (pLVX) backbone containing a CMV promoter to drive the expression of our S15-mCherry construct. Lentiviral particles were generated by PEI transfection of HEK293T cells. The transfection was performed with equal nucleotide concentrations of VSV-g, ΔNRF or psPax2, and pLVX-CMV-S15-mCherry plasmid over night and the cell medium was then changed. The cultured medium from the transfected HEK293T cells was collected 48 hours later and purified using a .45um filter and syringe (Millipore). The purified medium was then spinoculated onto the receiving HEK293T cells by centrifugation at 13°C for 2 hours at 1200 × g or directly added to THP-1. 72 hours later the cells were then selected for expression of our S15-mCherry construct by supplementing DMEM or RPMI 1640 with 5 ug/ml puromycin (Hyclone).

### Generation of GFP Fusion Protein Constructs

Expression plasmids containing HIV-1 proteins TAT, VPR, and NEF were all generated by PCR based cloning and restriction enzyme strategies. Primers against TAT, VPR, and NEF proteins were created and either first inserted into the pEGFP-N1 plasmid to create N-terminal GFP fusion constructs of TAT and NEF followed by subcloning into the lentiviral (pLVX) plasmid or directly inserted in pLVX C1-GFP in the case of VPR.

### Transfection of HIV1 GFP constructs

HEK293T cells were transfected at approximately 60 percent confluency using Polyethylenimine (PEI), and either pLVX TAT-GFP, GFP-VPR, or NEF-GFP plasmids over night. The media was changed and the transfected cells were cultured in media for 48hrs before the media was collected. The cultured media was concentrated via ultracentrifugation.

### Purification of Extracellular Vesicles

To purify extracellular vesicles, cultured media was collected from either HEK293T or THP-1 cells. The collected media was then sequentially centrifuged. 15ml or 50ml conical were first centrifugation in a tabletop centrifuge at 2000g for 20 minutes at 4C. The supernatant was then collected and added to either Beckman Coulter polycarbonate centrifuge tubes (#349622) or (#344058) and spun at 10,000g with either SW41 TI or SW28 Beckman rotors, respectively, in a Optima L-90K Ultracentrifuge at 4°C for 30 minutes. Subsequently, the supernatant was collected and then ultracentrifuged at 100,000*g* for 150 minutes at 4°C using new of the previously stated tubes, rotors, and ultracentrifuge. Afterwards, the supernatant was discarded and the pellet was resuspended in PBS. The resuspended pellet was then subjected to another round of 100,000*g* centrifugation with same rotor and machine for 150 minutes at 4°C. The supernatant was then discarded and the pellet was then resuspended overnight in 100ul of PBS on an orbital shaker.

### Western blotting & Western Antibodies

Non-reducing western blots conducted from pellets of cells or concentrated cultured media in lysis buffer composed of 100mM Tris pH 8.0, 1% NP-40, and 150mM NaCl and protease inhibitor cocktail (Roche) on ice for 30 minutes. The lysates were then centrifuged for 10 minutes at 10,000*g* and afterwards the supernatant collected. The collected supernatants’ protein concentrations were then determined by Pierce BCA protein assay kit (Thermo Scientific). An equal fraction of non-reducing SDS solution was then added to the proteins and the contents were boiled on a dry-block for 5 minutes. Subsequently, the protein contents were equally loaded and ran on a 10% polyacrylamide gel for SDS-polyacrylamide gel electrophoresis (SDS-PAGE). After separation, the proteins were transferred to a nitrocellulose membrane (Bio-Rad), and then probed overnight at 4°C with the respective primary antibodies diluted in powdered milk block solution at 2.5g/50mL of TBST: mouse anti-CD9 (BD Pharmigen #555370) 1:1000; mouse anti-CD63 (BD Pharmigen #5556019) 1:1000; mouse anti-CD81 (BD Pharmigen #555675) 1:1000; mouse anti-GAPDH (Santa Cruz SC-32233) 1:3000; or rabbit anti-mCherry (Novus bio NBP2-25157) 1:1000. The nitrocellulose was then washed in TBST and probed with the respective HRP conjugated donkey anti-mouse or anti-rabbit (ThermoScientific) diluted in milk block solution at 1:10,000 for 30 minutes. HRP was then detected with the addition of SuperSignal West Femto Chemiluminescent Substrate (Thermo Scientific) as measured by chemiluminescence levels using the FlourchemE Imaging System (Protein Simple).

### Transmission Electron Microscopy

Concentrated EVs were resuspended in PBS and 50ul of the total resuspension was then added to an equal volume of 4% paraformaldehyde. 5uL of fixed EVs were then added to a grid and allowed to incubate for 20 minutes.The grid was then washed twice by floating it 100ul PBS 3 minutes at a time.The grid was then floated in 4 separate aliquots of 100ul of 50mM glycine in PBS for 3 minutes. Afterwards, the grid was blocked by floating it in 100ul of block solution made up of 5% Bovine Solution Albumin (BSA) added to PBS for 10 minutes. The grid was then incubated with mouse anti-CD63 (BD Pharmigen #5556019) 1:50; mouse anti-CD81 (BD Pharmigen #555675) 1:50 which was diluted in 1% BSA PBS for 30 minutes by flotation. Subsequently, the grid was floated 6 times to wash it, each time in 100ul of PBS with .1% BSA for 3 minutes. The grid was then probed with donkey anti-mouse antibody conjugated to 20nm gold particles 1:200 (Cytodiagnostics #AC-20-02), diluted in block solution for 30 minutes by flotation. After, the grid was washed by floating it in 100ul of PBS with .5% BSA 6 times for 3 minutes each. The grid was then floated in 50ul of .1% glutaraldehyde for 5 minutes. It was then subsequently floated in 100ul of DI water 8 times at 2-minute intervals. Afterwards, the grid was floated in 50ul uranyl oxalate for 5 minutes, followed by 10 minutes of 50ul of methyl cellulose-uranyl acetate (MC-UA) on ice, and then blotted on filter paper to remove any excess. The grid was then allowed to dry for 10 minutes before being added to a metal box for extended drying over night before imaging.

### ImmunofluorescenceStaining

In order to adhere EVs on to coverslips, either 80uL of resuspended concentrated EV was added to 420ul of PBS totaling 500ul or, for unconcentrated EVs, 500uL of supernatant cultured media was added into the well of 24-well plate containing a glass coverslip. The contents of the 24-well plate were then spinoculated by centrifugation at 13°C for 2 hours at 1200 × g onto the coverslips and subsequently fixed in a solution of 0.1 M PIPES with 3.7% formaldehyde (Polysciences) for 15 minutes and washed 3x with PBS. The coverslips were then permeabilized with a .1% solution of saponin in block solution composed of 500mL of PBS supplemented with 10% normal donkey serum (NDS), and 0.01% NaN_3_ for 5 minutes. After washing 3x, the coverslips were then incubated with rabbit anti-Lamp1 antibodies (Abcam #24170) and either mouse anti-CD9 (BD Pharmigen #555370), mouse anti-CD63 (BD Pharmigen #5556019), or mouse anti-CD81 (BD Pharmigen #555675), in the previously stated block solution for 1 hour at room temperature. All primary antibodies were used at 1:1000. In experiments using lectins, biotin conjugated lectins(Vector Laboratories) were used at a working concentration of 5ug/mL in place of primary antibodies for 1 hour at room temperature. Afterwards the coverslips were washed with PBS and subsequently incubated with secondary antibodies of conjugated donkey anti-mouse 488 (Jackson ImmunoResearch Laboraties, Inc.) and donkey anti-rabbit 647 (Jackson ImmunoResearch Laboraties, Inc.) at a concentration of 1:400 for 30 minutes at room temperature diluted in PBS block solution and then washed with PBS. Additionally, FITC conjugated streptavidin (SAV) (Jackson ImmunoResearch Laboraties, Inc. 016-600-084) at 1:1000 was added for 1 hour at room temperature, diluted in PBS block, and then washed with PBS. Afterwards, coverslips were fixed and mounted.

### Human Bodily Fluid Sample Collection, Preparation, and Processing

#### Saliva

Whole saliva from a healthy donor was collected according to a previously published saliva collection protocol(24). Samples were stored at 4°C for no more than 36 hours, or at −20°C until needed. If frozen, samples were thawed at room temperature. Samples were then spun at 10,000 x *g* for 10 minutes to pellet cells and cellular debris. Due to the sticky nature of mucus-rich saliva, the supernatant from this first spin was transferred to a fresh tube and spun again at 12,000 x *g* for 15 minutes to remove residual mucus and cellular debris. The supernatant from this second spin was then passed through a 0.45µm filter into fresh tubes.

In these experiments, 200ul of saliva with mixed with 300ul of PBS so that a total volume of 500ul was added to glass coverslips in a 24-well plate. The plate was then spun at 13°C for 2 hours at 1200 x *g* to spinoculate the samples onto the coverslips. The coverslips were then fixed in a solution of 0.1 M PIPES with 3.7% formaldehyde (Polysciences) for 15 minutes and washed 3x with PBS, followed by a 5 min permeabilization step using a 0.1% saponin block solution composed of 500mL of PBS supplemented with 10% normal donkey serum (NDS), and 0.01% NaN_3_ and then washed another 3x with PBS. The spinoculated saliva was incubated with rabbit anti-Lamp1 antibody (Abcam #24170) and either mouse anti-CD63 (BD Pharmigen #5556019) or mouse anti-CD81 (BD Pharmigen #555675), along with biotin-conjugated wheat germ agglutinin (WGA) (Vector Laboratories, B-1025) at room temperature for 1 hour. All primary antibodies were used at a concentration of 1:1000 and all lectins were used at a final working concentration of 5µg/mL, diluted in a non-saponin block solution made up the same components of the previously stated solution, minus the detergent. Coverslips were then washed 3x with PBS, and incubated with secondary antibodies of conjugated donkey anti-mouse 594 (Jackson ImmunoResearch Laboraties, Inc.) and donkey anti-rabbit 647 (Jackson ImmunoResearch Laboraties, Inc.), and FITC conjugated streptavidin (SAV) (Jackson ImmunoResearch Laboraties, Inc. 016-600-084) at room temperature for 40 minutes. Antibodies were used at 1:400, and SAV was used at 1:2000, and all secondaries were diluted in the non-saponin block. Coverslips were then washed 3x with PBS and mounted onto to glass slides.

#### Plasma

Whole blood from a healthy donor was drawn into conicals containing 10%, by volume, of 3.8% sodium citrate. Lymphocyte separation medium (Corning 25-072-CV) was carefully layered into the bottom of the conical, and the whole blood was spun at 400*g* for 15 minutes. The plasma layer was carefully pipetted out, aliquoted, and immediately stored at −80°C until needed. Frozen samples were thawed at room temperature and then spun at 10,000*g* for 10 minutes to pellet cells and cellular debris. The resulting supernatant was then passed through a 0.45µm filter into fresh tubes.

In these experiments, 200ul of plasma with mixed with 300ul of PBS so that a total volume of 500ul was added to glass coverslips in a 24-well plate. Plasma samples were spinoculated, fixed, and stained in the same manner as the saliva samples detailed above.

### Wide-field Fluorescence Deconvolution Microscopy and Analysis

Images of the extracellular vesicles and cells were taken with the DeltaVision wide field fluorescent microscope (Applied Precision, GE) outfitted with a digital camera (CoolSNAP HQ; Photometrics), while using a 1.4 numerical aperture, and 60× objective lens and 100x objective lens respectively.

The amount of images taken per experiment is stated in the figure legend. In each case, the images taken were from different locations on the cover-slip with the intent of creating a representative sample of the total population. During some experiments, the “panels” function of the SoftWoRx software (Applied Precision) was used to take non-biased images with the same set of panels applied to each coverslip. In these cases, the z-stack distance was manually recalibrated for each coverslips to maintain that the images remained in focus. Images in each experiment were subjected to the same set of exposure conditions for each condition and replicate.The resulting collected images were deconvolved after their acquisition with the SoftWoRx deconvolution software (Applied Precision). The deconvolved images were then analyzed on Bitplane: Imaris software version 7.6.4, where the spots masking algorithm was built around the signal of interest (i.e. S15Ch, an HIV-fusion protein, or a lectin).

All S15 mCherry acquired images were subjected to the same spots or surface masking algorithm or for HIV1 proteins, via the Batch Coordinator tool (Bitplane) to each respective signal. Images in which the spots masking algorithm did not accurately capture the intended signal were thrown out of the sample population. No more than 2 images per condition per replicate were excluded.

Background levels of maximum intensity for each respective channel were determined based on secondary antibody controls for mouse (CD9, CD63, and CD81) or rabbit (LAMP1), and streptavidin(SAV)-only controls (WGA, LEL, etc) via a gating process analogous to flowcytometry. In cases where the threshold for above background signal was not visually obvious, the 99^th^ percentile was determined and used as the threshold.

### Statistical Analysis

All statistical analyses were conducted and graphs were made with GraphPad Prism version 6.00 (GraphPad Software, Inc.). Graphs with data from a single cultured media preparation and coverslip are shown demonstrating the mean with the standard deviation. Experiments in which multiple cultured media preparations and independent coverslips were imaged are shown with the mean and standard error of the mean. Two-tailed t-test with student’s post-hoc was used to compare data with a single independent variable and Two-way ANOVA with Tukey’s multiple comparisons test was used to conduct statistical test whether data comparing the differences in co-localization percentages for either differing analysis methods or cell-types was used. For all statistical tests: * = p-value < .05; ** = p-value < .01; *** = p-value < .001; and ****=p-value < .0001.

## Results

In order to allow EVs secreted from cultured cells to be visualized, we transduced HEK 293T cells to express a S15-mCherry construct (S15Ch). The S15Ch construct contains the 15 N-terminal amino acids of c-SRC appended to the N-terminal of the mCherry fluorophore sequence. It has previously been shown that these 15 amino acids are necessary and sufficient for the addition of the myristoyl lipid anchor on the N-terminal glycine of c-SRC, leading to its plasma membrane localization (25, 26). In experiments in which this construct was used to label HIV-1 viral particles(25, 26), we also observed that this construct is incorporated into EVs which lack HIV-1 p24 and is released from cells constitutively and independently of virus production, similar to the manner in which a palmitoylated fluorescent proteins are incorporated into EV populations when expressed in cells (27). To verify that this construct was incorporated into EVs, we collected culture supernatant from 293T cells expressing S15Ch following transient transfection and isolated EVs by differential ultracentrifugation. The S15Ch construct was observed within the ultracentrifuged EV fraction, which also contained the canonical exosomal proteins, CD9, CD63, and CD81(Fig 1A). To determine if the S15Ch protein was associated with EV membranes, we also added 0.1% SDS to cultured medium prior to ultracentrifugation. The addition of SDS, which causes disruption of lipid membranes, eliminated the presence of S15Ch recovered following ultracentrifugation (Fig 1A), demonstrating that S15Ch is incorporated into EVs. Moreover, when we subjected our concentrated EVs to transmission electron microscopy, we found that the concentrated fraction contained lipid enclosed structures between 30-150nm in diameter consistent with what is reported for EVs and were positive for CD63 and CD81(Fig 1B).

**Figure 1:**
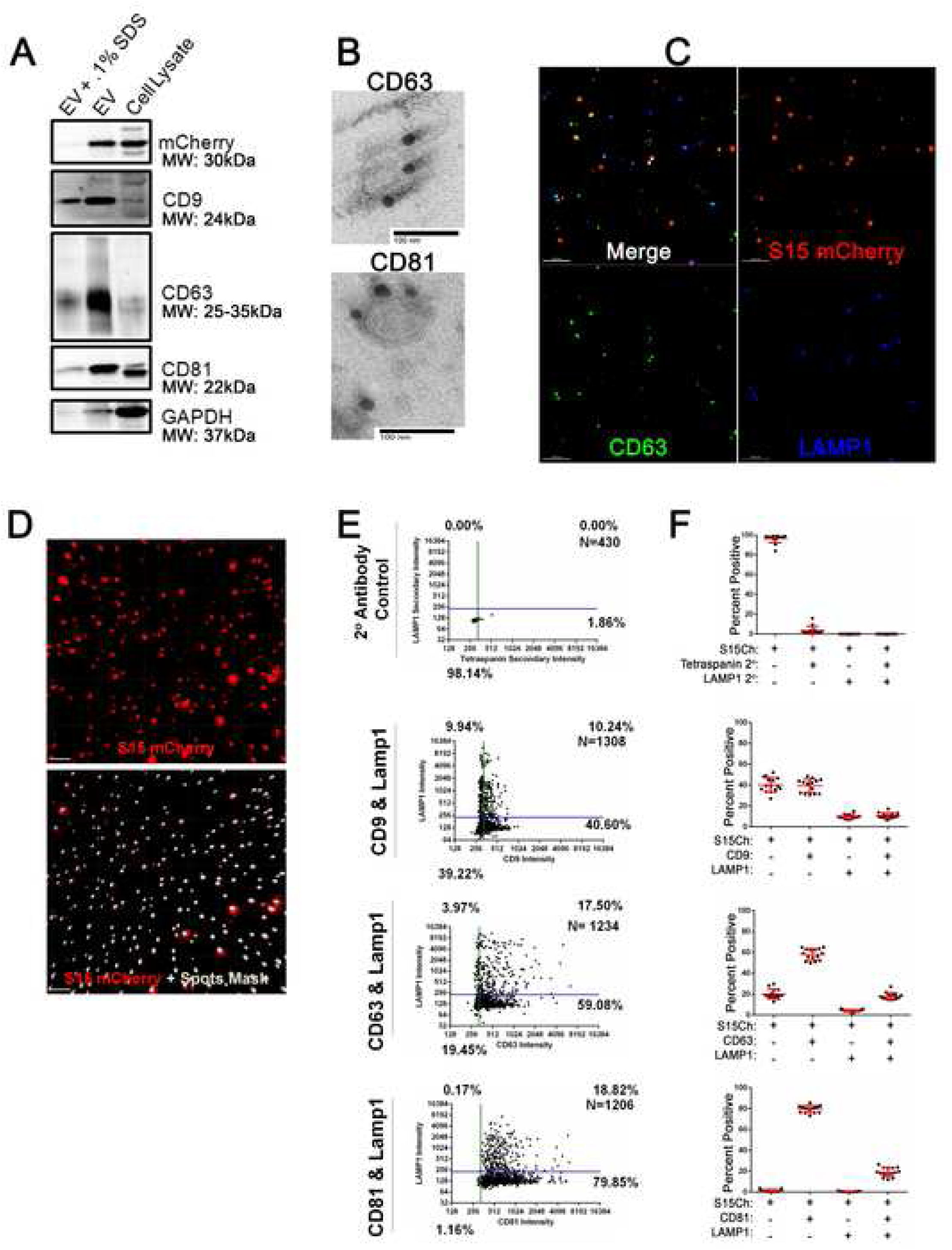
Detection of endogenous protein markers on EVs released from 293T cells. A. Non-reducing SDS-Page gel of mCherry positive 293T cell lysate or ultracentrifuge concentrated extracellular vesicles with or without the addition of SDS, probed with antibodies for mCherry, CD9, CD63, CD81, and GAPDH. B. Representative transmission electron microscopy images of concentrated S15 mCherry EVs showing primary antibodies against CD63 and CD81, respectively, and secondary anti-mouse conjugated to 20nm gold particles. C. Representative image from S15 mCherry HEK 293T cells’ cultured media spinoculated onto a coverslip. The individual mCherry, CD63, and LAMP1 channels are shown, respectively, in addition to the combined merge. D. The images show the S15 mCherry channel alone with the masking algorithm generated in Bitplane Imaris imaging software used to identify S15 mCherry signal. E. Masking algorithm in D was used to calculate the degree of EVs positive for the indicated protein. In secondary control, identical staining in the absence of primary antibody was performed. E. Data compiled from 20 images from a single coverslip. Percentage of S15 Spots described as positive or negative for each marker is indicated. Error bars display the mean and the standard deviation.

We next sought to determine if endogenous proteins which are known to be incorporated into EV populations could be detected on S15Ch EVs by immunofluorescence. To this end, we spinnoculated(28) EVs purified by ultracentrifugation or tissue culture supernatant filtered through a 0.45µm filter onto glass coverslips. This low speed centrifugation (2 hrs @ 1200*g*) is sufficient to significantly increase viral binding to cells and coverslips, allowing them to be interrogated by immunofluorescence following fixation (25, 28). Following fixation, EVs were stained with a mouse antibody against the tetraspanin protein CD63 and a rabbit polyclonal antibody directed against LAMP1, two proteins known to be present in EV populations. Following staining, coverslips were imaged using wide-field, fluorescent deconvolution microscopy, which is ideally suited for quantification of such specimens due to its sensitivity and field uniformity (29). Using this approach, a subset of S15Ch+ puncta could be observed to co-localize with both markers to some degree (Fig 1C), although not all CD63+ or LAMP1+ puncta were positive for S15Ch and not all S15Ch+ puncta were positive for CD63 or LAMP1.

### Characterization of EV populations by multiplexedimmunofluorescence imaging

We next determined if such an approach could allow for the reproducible multiplexed analysis of EV populations based on the markers present on EVs detected by indirect immunofluorescence. To this end, we stained individual coverslips with mouse antibodies to the tetraspanins CD63, CD81 and CD9 and a rabbit polyclonal antibody directed against LAMP1. Following data acquisition and deconvolution, software-based algorithms were generated to identify individual S15Ch puncta using defined size and intensity criteria and create individual masks around each punctumpresent in these images (Fig 1D). Following manual validation of the algorithm in a subset of images to ensure that individual puncta were reliably identified, the algorithm was applied to all images collected and the multiplexed intensity of tetraspanin and LAMP1 staining present in each S15Ch+ puncta was calculated for each image collected. Secondary antibody controls were used as a negative control to define the background staining intensity in each channel and establish threshold intensities above which individual EVs were described as positive for individual tetraspanins or LAMP1 staining (Fig 1E).

Multiplex analysis of EV populations identified by masking events identified in the S15Ch channel revealed differential staining of tetraspanin markers on these populations when data from individual images was examined. For example, over 98% of the S15Ch+ EVs analyzed exhibited CD81 staining above background, while approximately 50% were positive for CD9 (Fig 1E). In contrast, the amount of LAMP1 signal on these EVs was observed to be consistent, with approximately 20% of S15Ch+ EVs exhibiting LAMP1 staining above background (Fig 1E). Critically, when data from multiple images were compared as technical replicates, the percentage of EVs positive for one or both markers was observed to be extremely reproducible (Fig 1F). Additionally, we found that the mean values identified in this way were highly reproducible across multiple independent cultured media and coverslip preparations. Moreover, nearly identical outcomes were obtained if events identified in individual images were pooled or if data from individual images were considered individual replicates and averaged to determine the percentage of EVs positive for each marker (Fig 2A).

**Figure 2:**
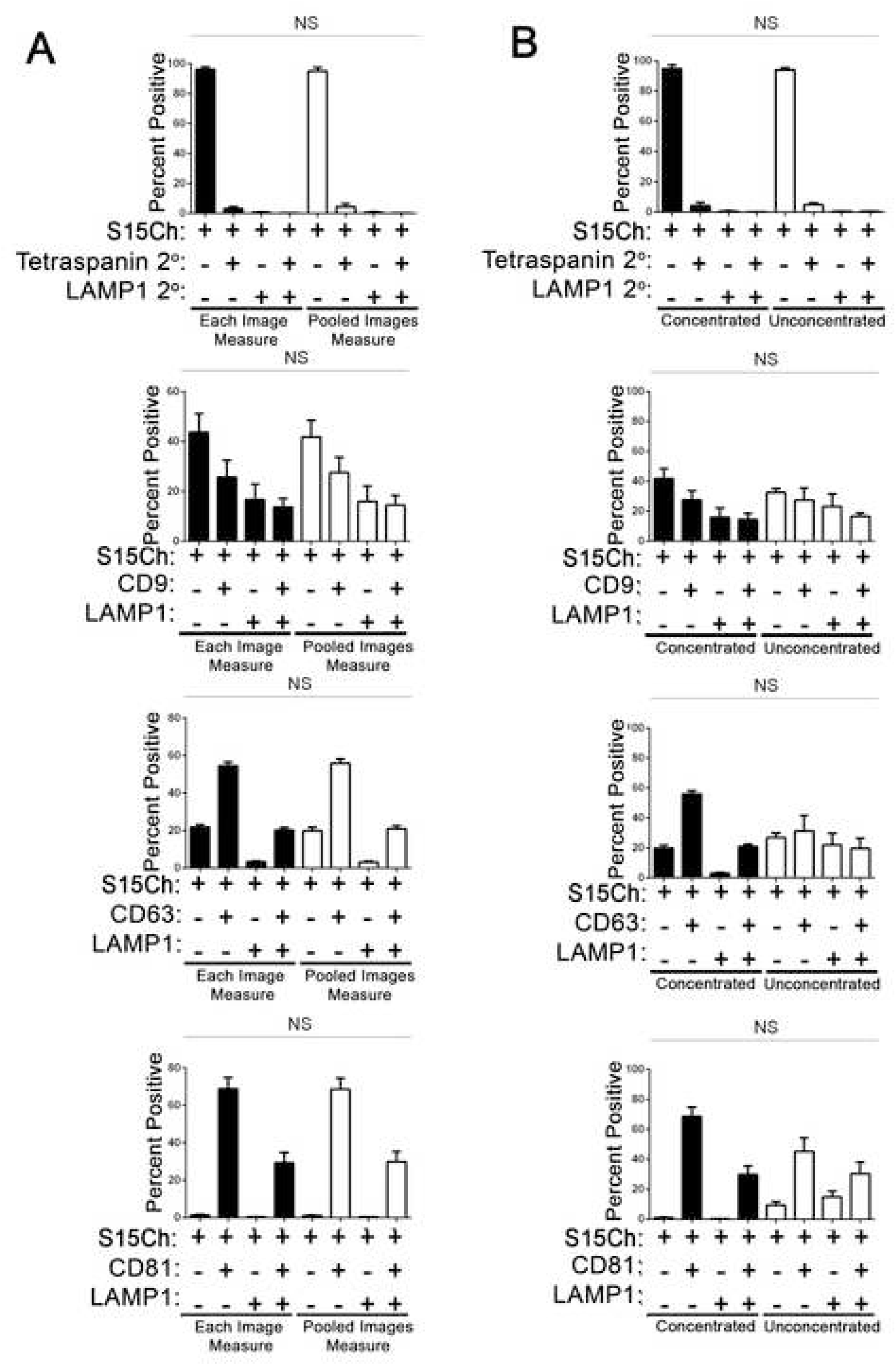
EV-MAC reproducibly stains EV populations following concentration or from tissue culture supernatant. A. Cultured media containing EVs were either left unconcentrated or concentrated via differential ultracentrifugation. The EVs were then stained with CD9, CD63, CD81, and LAMP1. Data shows the mean value from 3 separate coverslips from independently collected media in which 20 images were taken per coverslip. Error bars depict standard error of the mean. No significant differences were found among each respective staining paradigm among the unconcentrated and concentrated EVS when subjected to a -way ANOVA with Tukey’s multiple comparison post-hoc and found non-significant. B. Data compares the mean value among 3 independent experiments for S15 mCherry co-localization comparing the mean values of a single replicate as determined by either each images co-localization percentage or all the data from a single replicate pooled together. Each replicate was composed of 20 images. All data shown was subjected to two-way ANOVA with Tukey’s multiple comparison post-hoc and found non-significant. All graphs depict error bars showing the mean with standard error of the mean.

As we had previously observed EVs present in stocks of viral supernatant that had not been subjected to differential centrifugation, we next asked if we could detect differences between EV populations obtained through differential centrifugation and those spun directly onto glass from conditioned media. We did not detect a significant difference in the relative staining of tetraspanins or LAMP1 when EVs were enriched by ultracentrifugation or spun directly onto coverslips from conditioned media (Fig 2B). We did note that differential centrifugation tended to lead to the appearance of larger accumulations of signal that likely represent EVs aggregated during centrifugation. However, these aggregations could be excluded from the analysis by incorporation of size constraints into the mask algorithm (Supplemental Fig 1A) and generally reduced by thorough resuspension of the pelleted material (data not shown). These data demonstrate that multiplexed analysis of co-localization(EV-MAC) provides highly reproducible analysis of EV populations.

### EV-MAC reveals producer cell and cargo specific changes in EV populations

To determine if EVs released from different cells types exhibited a distinct staining profile, we collected EVs from a monocytic cell line, THP-1, stably expressing the S15Ch construct in parallel with our 293T cells. Pooled imaged analysis of S15Ch+ EVs released from our THP-1 cells revealed that this EV population appeared to be generally less positive for tetraspanins and more positive for LAMP1 (Fig 3A&3B,). Indeed, when we compared the S15Ch+ EVs from our S15Ch 293T and S15Ch THP-1 cell-lines, respectively, a smaller percent of THP-1 EVs were negative for LAMP1 and positive for CD9 or CD81and correspondingly, a larger percentage of the THP-1 EVs were positive for LAMP1 and negative for CD9 or CD81 (Fig 3C). A comparable percentage of the S15Ch+ EVs were triple positive for S15Ch, LAMP1, and either CD9 or CD63 but not for CD81 among both THP-1 and 293Ts. Finally, when we independently compared the total percent of S15Ch+ EVs positive for each respective marker, a significantly larger percentage of THP-1 EVs were positive for LAMP1 but a significantly lower percentage were positive for CD9 and CD81.These results demonstrate that EV-MAC can reliably and reproducibly identify differences in EV composition in EV populations from difference cell types.

**Figure 3:**
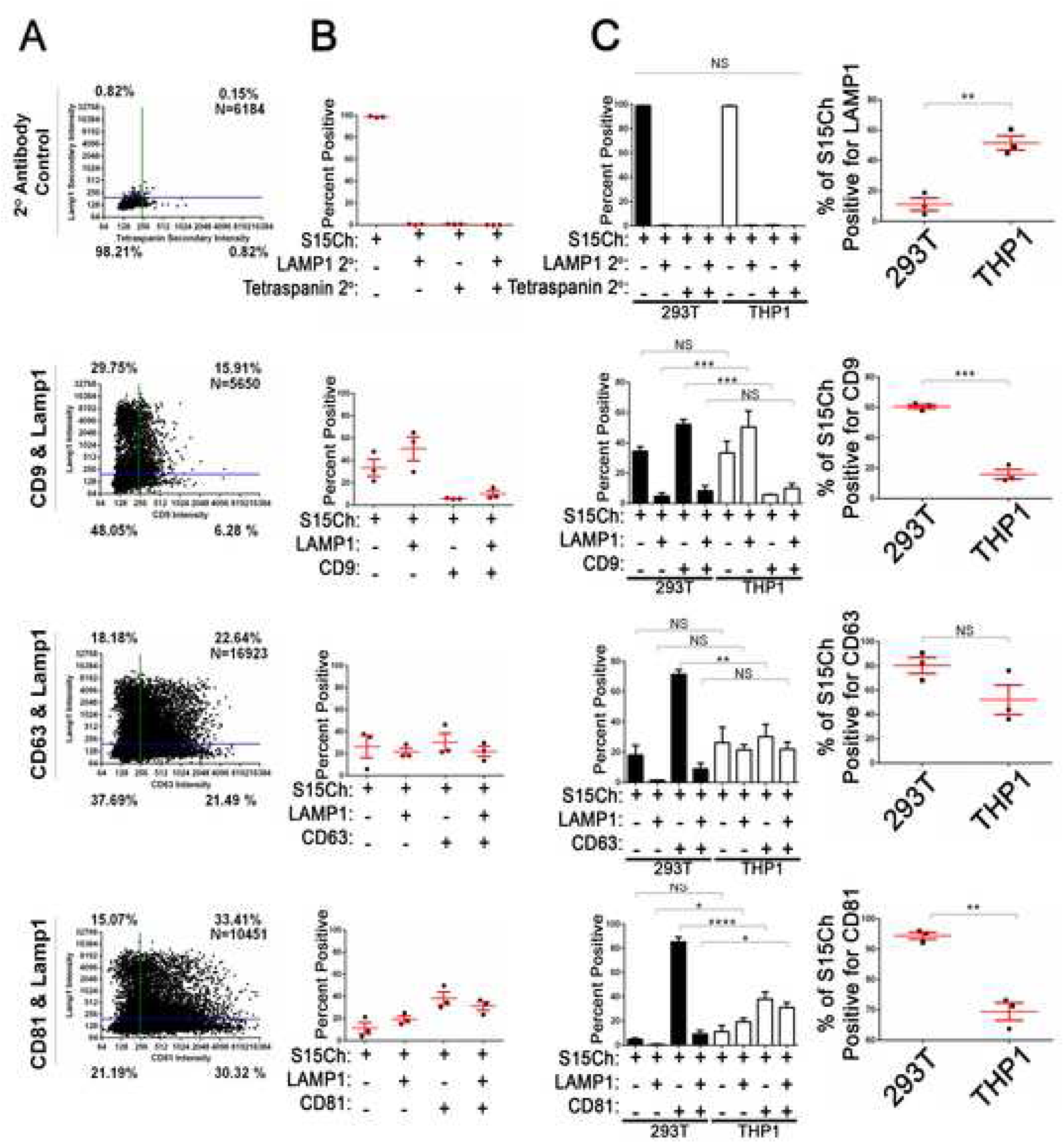
S15-mCherry THP-1 EVs co-localize with canonical EV markers at differing rates compared to 293T mCherry EVs. A.S15 positive EVs from THP-1s were stained for the indicated tetraspanin and LAMP1. Data from 15-20 images were pooled together from a single coverslip. B. The mean percentage of S15Ch+ EVs positive for each indicated marker from 3 independent media preparations and coverslips was calculated. Error Bars represent standard error of the mean. C) Comparison of tetraspanin and LAMP1 staining on EVs released from THP-1 and 293T cells. Left panel shows the data compared based on co-localization plot quadrants while the second set of graphs shows the total percent of EVs positive for each individual stain, respectively. Data shown compares the percent positive EVs collected from the cultured media of S15CH THP-1 and S15CH 293T cells which were then subjected to our staining paradigms. The first set of graphs the same masking algorithm was used to evaluate the S15Ch EVs from both cell lines. The data from the first column of graphs shown was subjected to two-way ANOVA with Tukey’s multiple comparison post-hoc. The data from the second column of graphs was subjected to Student’s Two-tailed T-test. All graphs shown depict mean value among 3 independent coverslips and media preparations and error bars show the standard error of the mean. * = p-value < .05, ** = p-value < .01, *** = p-value < .001.

We next asked if we could use EV-MAC to characterize EVs containing different cargo released from the same cell type. To this end, we utilized GFP-tagged versions of the HIV-1 proteins Tat, Vpr and Nef, as these proteins have all been implicated as cargo in potentially pathogenic extracellular vesicles released from HIV-1 infected cells (21). To determine if these cargos were released in EVs with discernable differences in canonical EV markers, we transfected GFP-tagged versions of Tat, Vpr and Nef into 293T cells and collected media from these cells for EV-MAC analysis. We examined the degree of CD81, CD63, CD9, LAMP1, as previously described, and also stained EVs for the ESCRT protein TSG101, which is known to be incorporated into EVs (30). For analysis, a masking algorithm was created around GFP+ signal and the degree of association of the markers of interest was determined. We observed that GFP-Neff+ puncta were positive for the tetraspanin markers examined, qualitatively similar to what was observed with S15ch+ puncta (Fig 4A). However, Nef-GFP+ puncta exhibited comparatively little LAMP1 staining compared to S15ch+ puncta. Approximately 10% of Nef-GFP+ puncta exhibited TSG101 staining above background (Fig 4A). By comparison, Tat-GFP+ puncta exhibited a greater degree of LAMP1 staining than Nef-GFP+ puncta, as were more likely to exhibit above background staining for TSG101 (Fig 4B), while a similar level of these EVs were positive for the tetraspanin markers examined. Finally, TSG101 staining was detected in a higher percentage of GFP-Vpr+ puncta compared to Tat-GFP and Nef-GFP puncta and CD63 staining was reduced in these puncta. These data demonstrate that EV-MAC can reveal differences in EV populations containing specific cargo.

**Figure 4:**
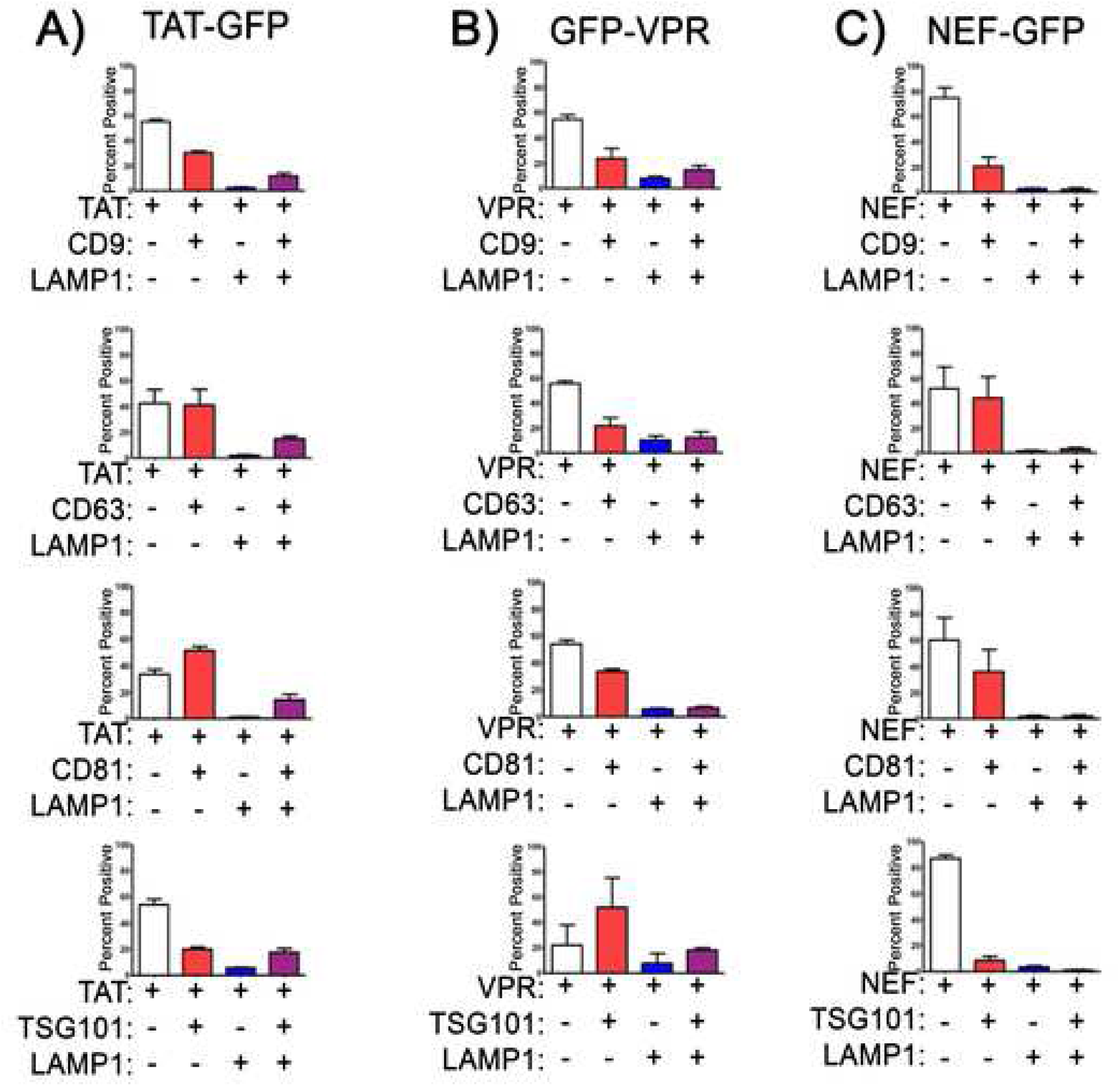
EVs containing different HIV-1 proteins exhibit differential tetraspanin and LAMP1 staining: The degree of co-localization of (A) TAT-GFP,(B) GFP-VPR, and (C) NEF-GFP EVs with tetraspanin markers, LAMP1 and TSG101. The data shown is the mean value from 3 independent media preparations and coverslips in which pooled data from a single coverslip was used to determine the mean co-localization percentages. Error bars show the standard error of the mean.

### EV-MAC identified changes in lectin binding in EVs released following lysosomal dysfunction

We also examined the glycan composition associated with S15ch+ EVs by staining with a panel of biotinylated lectins which bind to carbohydrates of differing sugar linkages present on glycoproteins and glycolipids present in EVs. Furthermore, we also assessed whether inhibition of lysosomal acidification altered the glycan composition by treating cells with bafilomycin-A1. Bafilomycin-A1 has previously been shown to alter the secretion of both EVs and cargoes that undergo autophagic driven secretion (13, 31). Among the lectins tested, we found that when lysosomes were not inhibited ∼85% or more of the S15Ch+ EVs were positive for *Lens culinaris* agglutinin (LCA); *Lycopersicon esculentum* (Tomato) Lectin (LEL); Pisumsativum agglutinin (PSA) and *Ricinus communis* agglutinin-1 (RCA1); *Erythrina cristagalli* lectin (ECL); *Phaseolus vulgaris leuco*-agglutinin (PHA-L); soybean agglutinin (SBA); and *Solanum tuberosum* (Potato) lectin (STL) were ∼50-85% positive; *Griffonia simplicifolia* lectin1&2 (GSL1) & (GSL2); peanut agglutinin (PNA); *Ulex europaeus* agglutinin 1 (UEA1); and *Vicia villosa* agglutinin (VVA) were ∼25-50%; and *Sophora japonica* agglutinin (SJA) as well as Succinylated Wheat Germ agglutinin (sWGA) were less than 25% positive (Fig 5). Interestingly, we found that bafilomycin-A1 treatment altered the lectin binding of S15Ch+ EVs, with significant reductions measured in ECL, LEL, PHA-L and PSA staining (Fig 5). Strikingly in contrast to other tested lectins, RCA1 and PNA co-localization did not follow this observed trend and remained unaffected. Collectively our data show that lectin binding to EVs can be assessed microscopically in the context of the EV-MAC workflow and reveals that lysosomal inhibition alters the glycan composition of EVs released under basal conditions, based on the observed changes in lectin binding profile.

**Figure 5:**
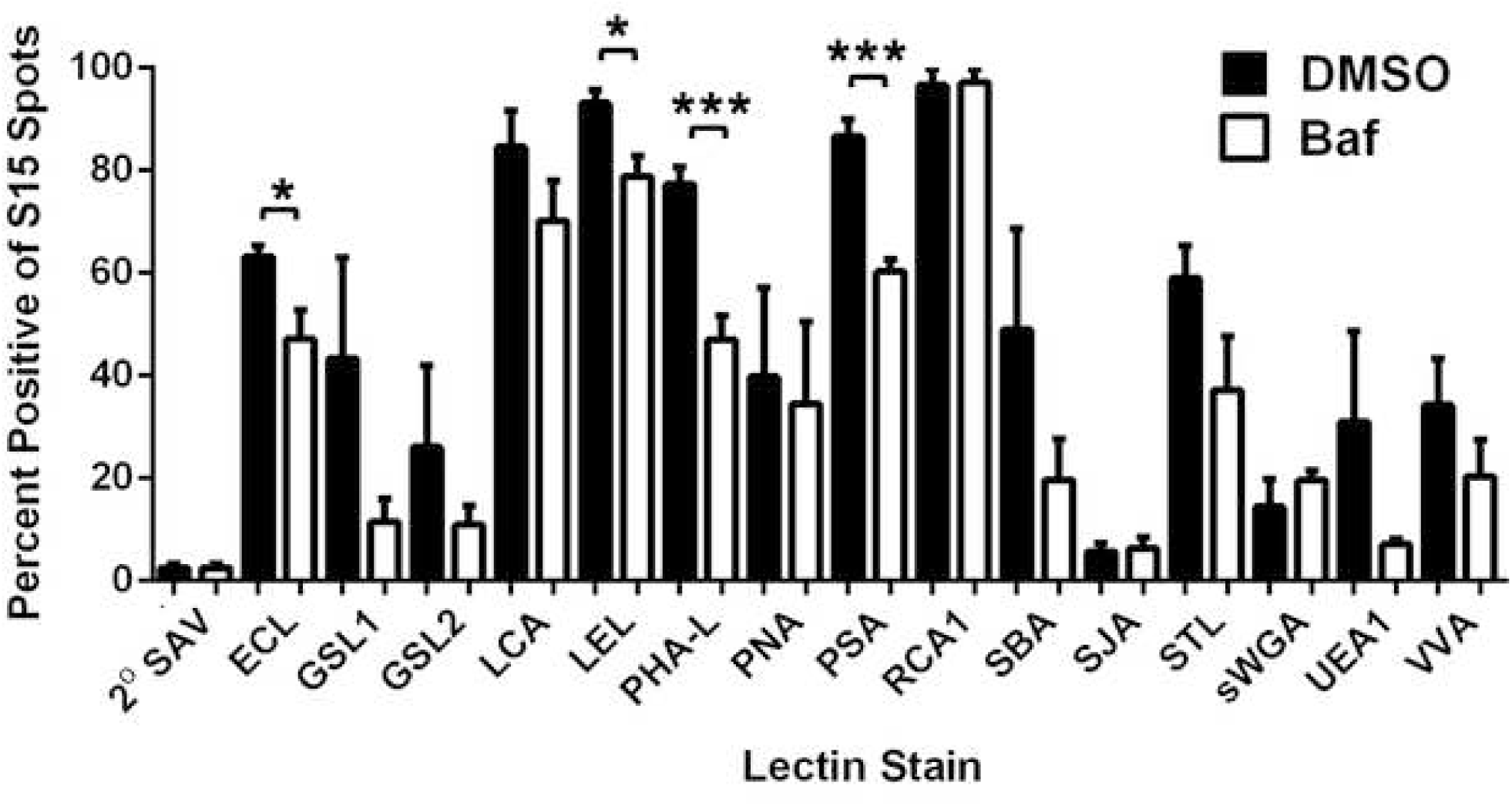
S15Ch+ EVs and Lectin co-localization is altered during lysosomal impairment. The cultured media from S15Ch 293Ts was incubated with the indicated biotinylated lectins followed by Alexa 488 streptavidin to determine the glycan profile of EVs under both DMSO Vehicle (DMSO) and 100nM Bafilomycin-A1 (BafA1) treated conditions. Data shown is the mean percent co-localization summation of at least 3 independent media preparations and coverslips. Significant differences between control and BafA1 treated samples were determined via two-tailed T-test. * = p-value < .05, ** = p-value < .01, *** = p-value < .001.Error bars show standard error of the mean.

### Application of EV-MAC to human biological samples

After validating this method with EVs collected from cell culture supernatants, we asked if our technique could be used to characterize EVs found in human bodily fluid samples. Samples of saliva and plasma were paneled with combinations of the lectin Wheat Germ agglutinin (WGA) and the LAMP1, CD63 and CD81 antibodies used in our cell culture studies. For analysis, masks were created around WGA+ signal and the degree of association of the tetraspanins and LAMP1 was determined.

In saliva, we observed approximately 15% of WGA+ puncta to be positive for CD63 and approximately 15% to be positive for CD81 (Fig 6). In general, less than 10% of WGA+ puncta were positive for both a tetraspanin and LAMP1. In plasma, we observed higher degrees of association of WGA with our markers of interest than in saliva. Approximately 17% of WGA+ puncta were positive for CD63, approximately 27% were positive for CD81, approximately 16% were positive for both CD63 and LAMP1, and approximately 17% were positive for both CD81 and LAMP1 (Fig 6). These data demonstrate that EV-MAC can be used with biological samples, and can reveal differences in the EVs obtained from different biological fluids.

**Fig 6:**
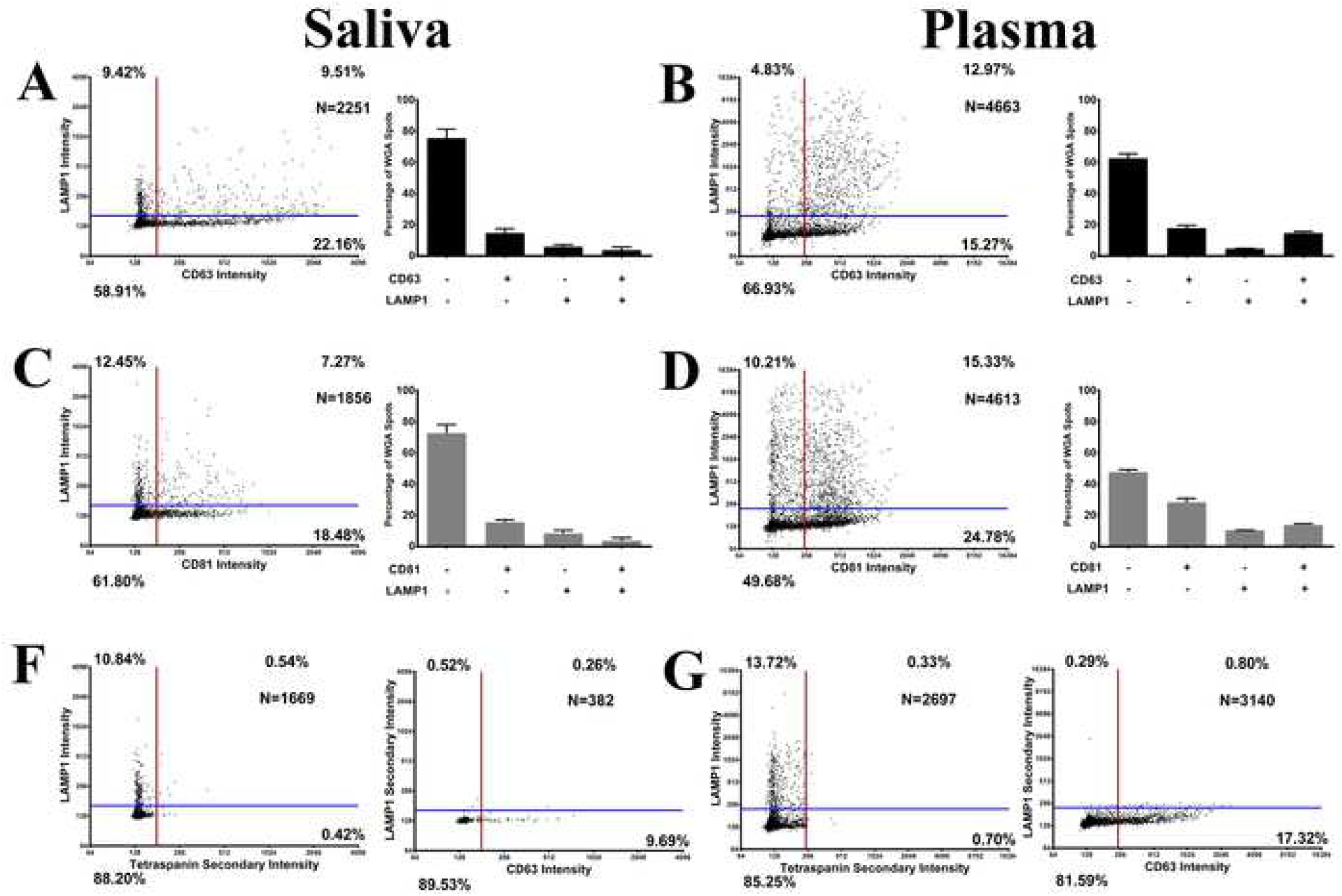
EV-MAC of EVs from Saliva and Plasma: EVs saliva and plasma were bound to glass coverslips via spinocculation and stained with antibodies to tetraspanins, LAMP1 and SAV. SAV+ puncta were evaluated for the presence of CD63 and LAMP1 (A,B) or CD81 and LAMP1 (C,D). Left panels show data compiled from individual coverslips and right panels show average of 3 or more technical replicates of the same sample. Error bars represent SEM. (F,G) Secondary antibodies controls for SAV+ puncta from each fluid.

## Discussion

Prior studies of extracellular vesicles have implicated EVs and their putative cargoes in a wide variety of cellular processes and pathological conditions. However, it is also increasingly clear that cells release a wide variety of EVs which vary in their cargo, membrane composition and biogenesis(8-10, 18, 19). While another study used a similar approach to assess co-localization of fluorescent markers incorporated into EVs (27), we demonstrate here that endogenous protein and glycan markers on EVs can be measured and used to characterize EV populations released from cells in culture or biological fluids. This was accomplished by using a workflow that we havetermed EV-MAC, which allows multiplexed analysis of EV populations according to their incorporation of canonical EV markersincluding tetraspanins, TSG101 and LAMP1. We first validated this workflow by utilizing the S15Ch+ EVs released from transduced cells by creating masking algorithms around the S15Ch signal to determine relative co-localization of our other markers of interest. Additionally, the generation of this masking algorithm allowed us to analyze images in a semi-high throughput manner and evaluate thousands of EVs in a short period of time with a small quantity of sample. The reproducibility of this approach across biological and technical replicates (Fig 2, Fig 3) demonstrates that this technique has value in its ability to comparatively analyze EV populations released from cells or present in biological fluid in normal and pathological states. As is the case with flow cytometry, the degree of EVs positive for any given marker will depend on the amount of the target antigen present in the vesicle and the specificity of the antibody being used, such that it would not be appropriate to suggest that a given protein is completely absent from an EV or population of EVs. However, as is also the case with flow cytometry, relative differences in staining between samples are likely to be very reliable and reproducible when a large number of events are assays, as demonstrated in this study.

Our observation that changes in the tetraspanins and LAMP1 content occurs in a cargo specific manner in EVs containing different HIV-1 proteins suggests that EV-MAC may be particularly useful in characterizing exosomes with pathological cargo, such as viral proteins or amyloid proteins associated with neurodegenerative disease or other situations in which EVs containing a specific cargo may represent a small percentage of total EVs in a sample. As such, this workflow may be useful in the identification of EV biomarkers that might not be identified when populations of EVs are measured *en mass* via proteomics or western blot.

To this end, we also performed an assessment of lectin binding to EV populations. While we did observed that lysosomal inhibition altered the binding of some lectins to EVs released from THP-1 cells, we also identified lectins which did not change under such conditions and found that ∼100% of EVs were identified by RCA1 staining. While the differential binding of lectins may be useful to understand the biogenesis of some EV populations and understanding the cell biology of the EV releasing cell (32-34), the use of lectins to broadly identify EVs to allow EV-MAC analysis of biological samples, especially in cases where the amount of sample is small and finite. In proof of this principle, we performed EV-MAC analysis on saliva and plasma samples, using lectin binding to create masks for subsequent analysis, similar to the role S15Ch played in other experiments.

Collectively, this manuscript established a workflow to monitor EV heterogeneity and identify cell type and cargo specific markers associated with individual EVs in a population. This workflow is likely amenable to other fluorescent microscopy methods and post-acquisition analytical methods, and thus provides an opportunity to better understand the cell biology of EV biogenesis under basal and pathological conditions and to define the composition of pathological EVs in biological fluids, especially when such pathological EVs represent a small fraction of EVs in a sample.

---

**Supplemental Fig 1: Masking Algorithms selectively removes S15Ch+ Aggregates.** Representative image of the S15Ch+ EV spots masking algorithm and its ability to selectively remove large aggregated S15Ch+ puncta and only include smaller punctum.

